# Saturated cell lysing is critical for high sensitivity microbiome analysis

**DOI:** 10.1101/2022.06.04.494821

**Authors:** Ying Wang, Cuiping Zhao, Yan Y. Lam, Liping Zhao, Guojun Wu

## Abstract

For robust DNA-based gut microbiome analysis, all cells in the stool samples need to be lysed. However, no standards have been developed to evaluate a DNA extraction protocol’s capability of lysing all cells and its sensitivity on detecting microbial structural differences among samples. In this study, we incrementally increased the intensity of mechanical lysis and integrated lysozyme pretreatment to Protocol Q (PQ), which was recommended as the best from 21 protocols^1^. A new protocol (LPQ) was optimized when DNA yield, Gram-positive bacteria ratio, and beta diversity all reached to a plateau with no further significant changes, indicating the achievement of saturated lysing. LPQ detected significant differences among three groups of fiber-treated human stool samples and identified 64 responsive ASVs, while a commercial kit failed to detect any significant treatment effects and PQ only detected 17 responsive ASVs. Therefore, saturated lysing as defined in this study should be adopted for evaluating microbiome DNA extraction protocols.

## Main Text

Gut microbiome is a complicated community consisting of approximately 3.8 × 10^13^ bacteria cells in colon^2^. These bacteria are belonged to around 500-1000 species with widely different cell wall structures^3^. For example, Gram-negative bacterial cell walls contain less peptidoglycan (<10%)^4^, a polymer that is responsible for rigidity of the cells, which makes them more vulnerable to lysing. Gram-positive bacterial cell walls contain more peptidoglycan (30-70%)^4^, which make them more resistant to lysing. If the cell lysing condition in a DNA extraction method is not rigorous enough, some cells in the samples may stay un-lysed which will introduce bias on microbiome analysis, e.g., leaving some critical members of the microbiome undetected.

So far, few studies have investigated whether a DNA extraction protocol has rigorous enough lysing condition that ensure complete lysing of all different types of cells in a microbiome sample to minimize bias in microbiome analysis. Two studies found that there was a positive correlation between prolonged mechanical lysing time with the DNA yield and the estimated sample diversity calculated from the sequential analysis^5, 6^. However, the microbiome analysis methods they applied were Temperature Gradient Gel Electrophoresis Analysis (TGGE) and Terminal restriction fragment length polymorphism (TRFLP) which were of low resolution. More recent studies on DNA extraction methods focused more on evaluating pre-existing commercial kits or protocols with next generation sequencing analysis. Schiebelhut et al. compared 6 pre-existing methods on DNA yield, DNA purity, extraction efficacy and cost and recommended using CTAB phenol–chloroform method^7^. In another study, Protocol Q (PQ) was considered the best after 21 pre-existing DNA extraction protocols were compared on Gram-positive bacteria ratio, protocol extraction accuracy based on mock community samples, and reproducibility^1^. Taken together, there is a lack of quality control standard for ensuring complete cell lysing by a DNA extraction protocol used for microbiome analysis.

To estimate whether the cell lysing condition in PQ has enough rigor for fecal samples analysis, we first incrementally increased the number of 8 min 15 s bead-beating from 2 times to 5 times on fecal samples collected from 4 self-claimed healthy young subjects with normal BMI and different ethnicities (Supplementary Table 1). Next, we applied 16S rRNA gene v4 region sequencing on the DNAs extracted under different bead-beating cycles and analyzed the data at amplicon sequence variant (ASVs) level, which provides the resolution down to the level of single-nucleotide differences^8^. We aimed to find out the optimized condition when DNA yield, beta diversity and Gram-positive bacteria ratio stop to increase significantly while the lysing rigor continue to increase. We hypothesize that when these three parameters reach a plateau, the complete/saturated lysing of different cell types in the samples will be achieved.

When we increased the number of bead-beating from 2 (PQ) to 5 times, DNA yield showed an increased trend (Fig. 1a). DNA yield in 2-Beatings group (PQ, 234 ± 16.5 ng/mg feces) was significantly lower than all the other groups. This indicated that more cells were lysed when increasing the intensity of lysing condition of original PQ. DNA yield in 3-Beatings group (301 ± 16.7 ng/mg feces) was significantly lower than that in 5-Beatings group (368 ±21.6 ng/mg feces). But no significant difference in DNA yield between 3-Beatings and 4-Beatings (342 ± 18.0 ng/mg feces) and between groups 4-Beatings and 5-Beatings was found. This indicated that DNA yield increased when increasing the number of bead-beating time and the plateau was reached after 4 times of bead-beating.

**Fig. 1.**
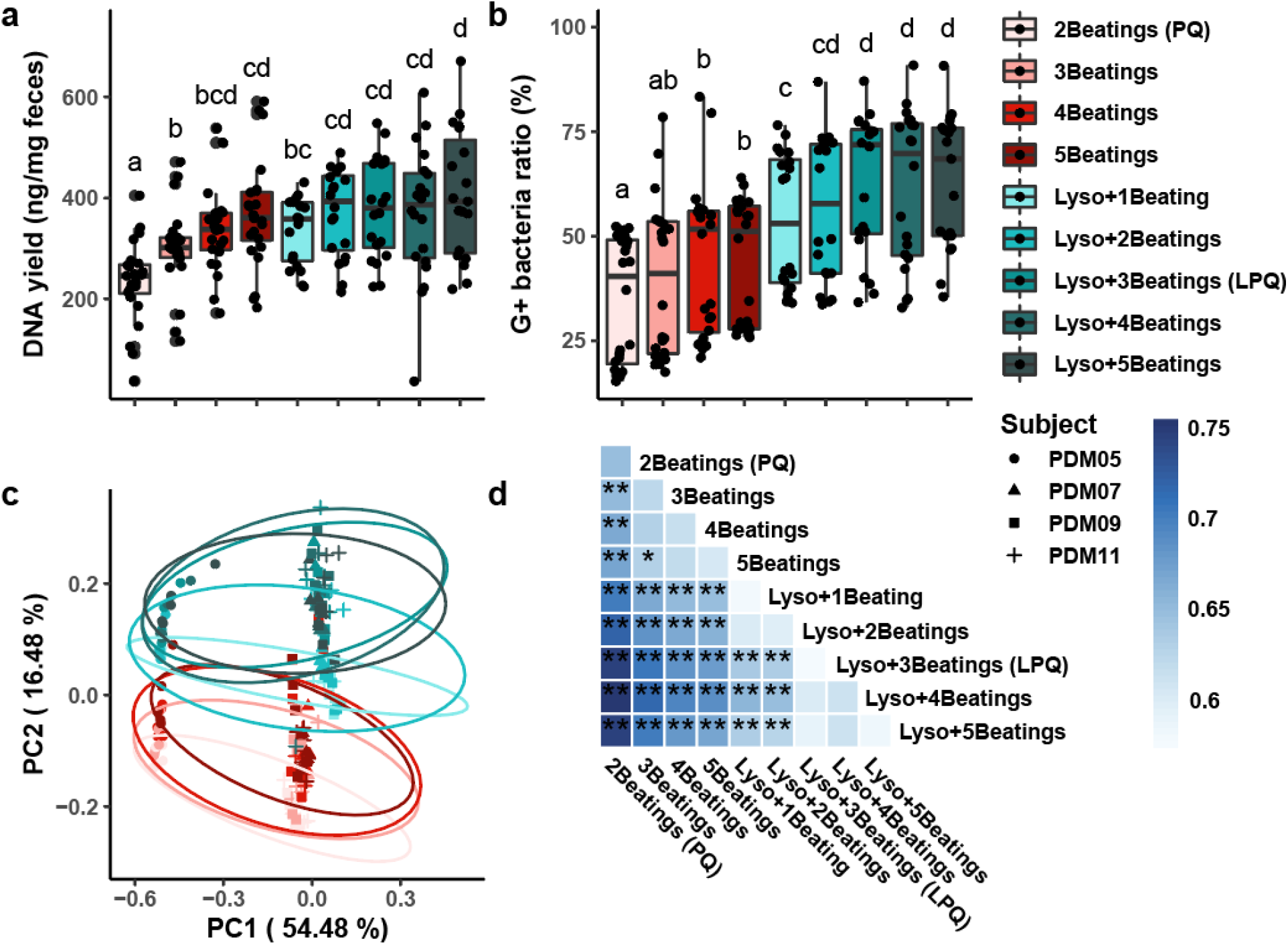
Combination of mechanical lysing with lysozyme pretreatment significantly increased DNA yield, G+ bacteria ratio and detected more diversity difference from human fecal samples. **a**, DNA yield. **b**, G+ bacteria ratio. Linear mixed-effects model with subject as random effect and Tukey’s test was used in **a** and **b**. Adjusted p value < 0.05 between two groups was considered significant and shown as different letters. **c**, Covariate Adjusted Principal Coordinates Analysis (aPCoA) with subject adjusted based on Bray Curtis distance derived from absolute abundance of the bacteria was performed on all samples. 95% confidence ellipses were shown with corresponding colors. **d**, Permutational multivariate analysis of variance (PERMANOVA) with subject adjusted was performed on Bray Curtis distance between groups and shown as asterisks (*P < 0.05, **P < 0.01). Mean Bray Curtis distance among samples between and within groups were shown in shades of blue. N= 4 fecal samples from 4 healthy participants were processed with modified lysing conditions with 5-6 replicates.

The ratio of Gram-positive bacteria also significantly increased when bead-beating times increased to 4 times (44.6 ± 3.7 %) and 5 times (43.8 ± 3.0 %) compared with 2-Beatings (34.7 ± 3.1 %) (Fig.1b). There was no significant difference in Gram-positive bacteria ratio between groups 3-Beatings, 4-Beatings and 5-Beatings indicating that Gram-positive bacteria ratio reached plateau after 3 times of bead-beating.

Weighted abundance of each ASV was calculated by multiplying the relative abundance of the ASV and the DNA yield per mg feces from DNA extraction. Bray Curtis distance was then calculated based on the weighted abundance of ASVs. Microbiome composition based on Bray-Curtis distance showed significant changes between samples extracted using 2-Beatings and all the other groups (Fig. 1c, d). Although no significant difference was found between 3-Beatings and 4-Beatings, and 4-Beatings and 5-Beatings, 3-Beatings showed significantly different beta diversity compared with 5-Beatings. This indicated that the plateau might be reached after 4 times of bead-beating.

Taken together, the lysing condition of PQ (2-Beatings) might not be enough to achieve complete lysis. 4-Beating group reached the plateau since no significant difference was found between 4-Beating and 5-Beating group. More bead-beating than 5 times was not attempted because of the time and labor consumption. To achieve more complete cell lysis in more efficient way, we decided to use lysozyme to pretreat the samples so that the glycosidic bond in peptidoglycan of cell walls for Gram-positive bacteria may be partially degraded and become more amenable to mechanical lysing^9^.

Single round of bead-beating combined with lysozyme pretreatment (Lyso+1-beating) can yield significantly higher quantity of DNA compared with PQ (Fig. 1a). No significant difference on DNA yield was detected between groups using 2 to 5 times of bead-beating combined with lysozyme pretreatment, indicating DNA yield reached plateau after 2 times of bead-beating combined with lysozyme pretreatment (Fig. 1a).

Similar to the results of DNA yield, Lyso+1-beating recovered significantly higher Gram-positive bacteria than all groups without lysozyme pretreatment emphasizing the efficiency of lysozyme on lysing Gram-positive bacteria (Fig. 1b). There was no significant difference found in Gram-positive bacteria ratio between groups Lyso+2-Beatings, Lyso+3-Beatings, Lyso+4-Beatings and Lyso+5-Beatings. This indicated that the Gram-positive bacteria ratio reached plateau after 2 times of bead-beating with lysozyme pretreatment as well.

Principal coordinate analysis (PCoA) showed that groups with more times of bead beating combined with lysozyme pretreatment have higher PC2 values which was consisted with samples processed without lysozyme pretreatment (Fig. 1c). This consistent trend indicated that the microbiome structure of a fecal sample changed along one direction, PC2 in this study, with the increase of intensity of lysing condition. There was no significant difference found between Lyso+3-Beatings, Lyso+4-Beatings and Lyso+5-Beatings (Fig. 1d). This indicated that beta diversity of samples reached the plateau after 3 times of bead-beating with lysozyme pretreatment.

All the above results showed that lysozyme pretreatment with more than 3 times of bead-beating achieved plateau in DNA yield, beta diversity, and Gram-positive bacteria ratio. Therefore, lysozyme pretreatment with 3 times of bead-beating (LPQ) was considered the protocol that can achieve saturated cell lysing.

To further investigate the effect of different protocols on microbiome analysis, we used three methods including Qiagen PowerFecal commercial kit (PF), PQ (2-Beatings), and LPQ to extract DNA from a same set of samples (N = 45) collected from an interventional study. In this study, 5 type 2 diabetes mellitus (T2DM) patients were recruited to donate one fresh fecal sample for an in vitro fiber fermentation study. Each fecal sample was homogenized in PBS buffer, filtered with cheese cloth, and cultured with three different dietary fiber groups including 1% bran fiber (BF), 1% inulin (IN) and 1% inulin plus 1% bran fiber (IN+BF) for 12 hours with triplicates. After 12-hour incubation under 37°C, three aliquots were collected from each sample and processed with the three DNA extraction methods respectively (N = 135). DNA yield of samples extracted using LPQ was significantly higher than that using PF in all three treatment groups and were significantly higher than PQ in IN treatment group (Fig. 2a). This indicated that LPQ lysed more cells and retrieved more DNA in different samples compared with both PF and PQ. Gram-positive bacteria ratio in LPQ was found significantly higher than both PF and PQ in all three treatment groups and PQ was only found significantly higher than PF in BF treatment group (Fig. 2b). This indicated that LPQ with the lysozyme pretreatment increased the Gram-positive bacteria lysing consistently from all the samples. Bray Curtis distance between samples was calculated and then visualized by Covariate adjusted principal coordinates analysis (aPCoA) with subject as the covariate to be adjusted (Fig. 2c). Samples from the same extraction method were clustered together indicating the method effect on microbiome analysis. Samples from different treatments using the same DNA extraction method were compared pair wisely on gut microbiome composition based on Bray-Curtis distance using subject stratified PERMANOVA test (Fig. 2d). No significant difference between treatments was detected using PF. Significant difference was detected when comparing treatments BF vs IN (adjusted p = 0.03) and treatments BF vs IN+BF (adjusted p = 0.03) using PQ. More significant difference (BF vs IN adjusted p = 0.0075, BF vs IN+BF adjusted p = 0.0075) in above two comparisons as well as marginally significant difference between IN vs IN+BF (adjusted p = 0.089) were detected using LPQ. To identify the ASVs that contribute to the significant difference between the treatment groups, prevalent ASVs (>20%) were compared between treatments using MaAsLin2 with subject as a random effect^10^. 17 ASVs were detected significantly different between treatments using PQ and 64 ASVs were found significantly different between treatments using LPQ (Figure 2c, d). Among all detected ASVs, 15 were identified by both PQ and LPQ, indicating that LPQ detected more difference compared with PQ. All the above results indicated that LPQ is the most sensitive in detecting treatment effects among the three DNA extraction methods.

**Fig. 2.**
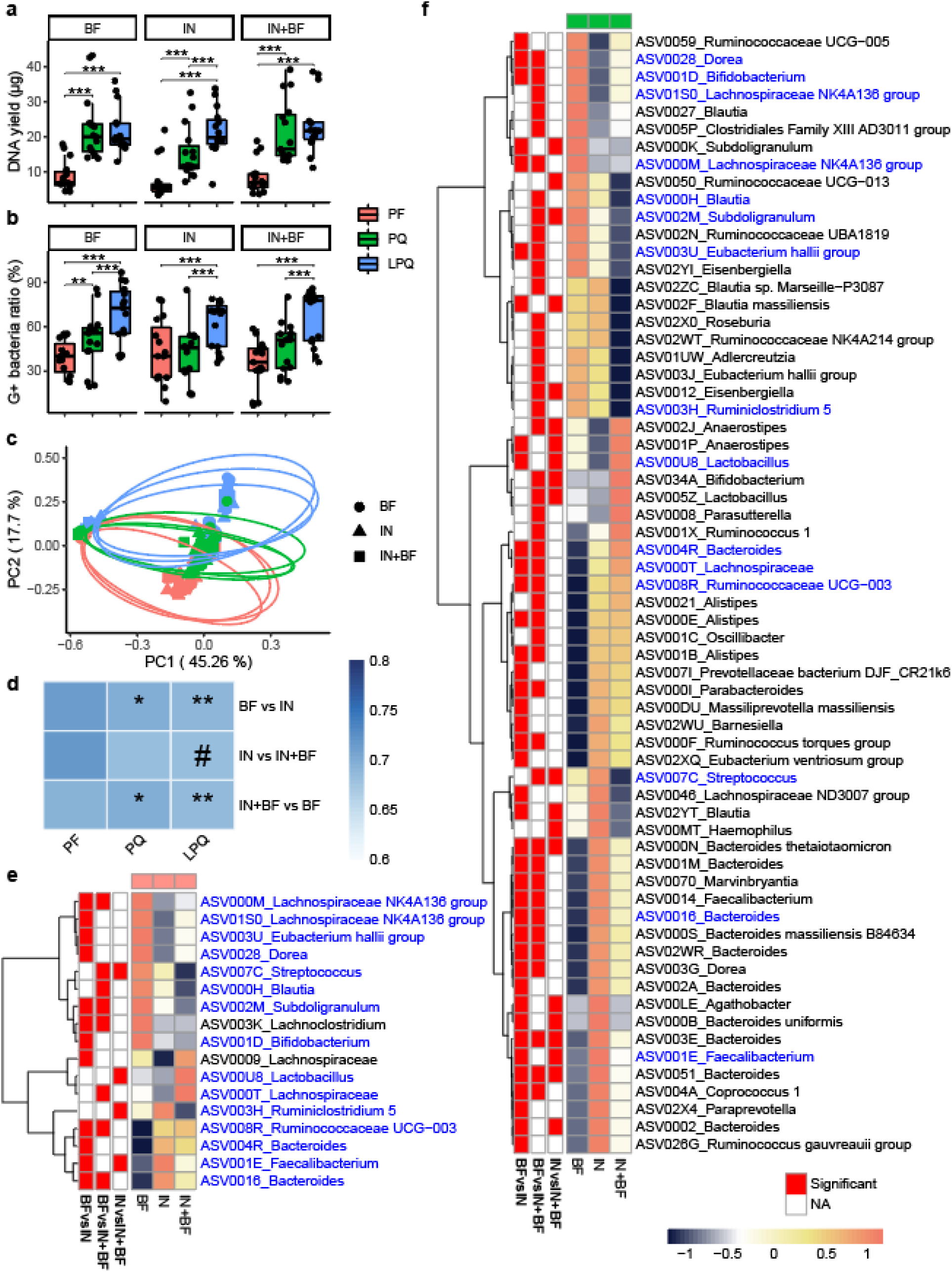
LPQ generated higher DNA yield and G+ bacteria ratio and was more sensitive to detect treatment effect. **a**, DNA yield and **b**, G+ bacteria ratio among samples from 5 T2DM patients with 3 treatments and different lysing conditions were shown. Linear mixed-effects model with subject as random effect and Tukey’s test was used and the significance was shown as asterisks (*** P < 0.001, ** P < 0.01, * P < 0.05). **c**, Covariate Adjusted Principal Coordinates Analysis (aPCoA) based on Bray Curtis distance were performed. 95% confidence ellipses were shown with corresponding colors. **d**, Statistical analysis on the beta diversity distance. Permutational multivariate analysis of variance (PERMANOVA) was performed between the three groups with subject adjusted and shown as asterisks (** P < 0.01, * P < 0.05, # P < 0.1). Mean distance between groups based on Bray Curtis distance was shown in the heatmap. **e**, 17 ASVs from PQ group and **f**, 64 ASVs from LPQ group were detected significantly different between 3 fiber treatment groups. 15 ASVs detected using both protocols were marked in blue. Three treatment groups were compared pair wisely for each extraction method. ASVs that were found significantly changed (adjusted p value < 0.25) were shown as red in the annotation heatmap on the left. Mean relative abundance for each group (N=5 subjects, 3 replicates per subject) was calculated and scaled for each row to generate the heatmap. ASVs were clustered using ward.D2-linkage method and is shown on the left.

In summary, our results showed that protocol Q which was recommended comparing between 21 representative protocols is still far away from saturated lysing all cells in all stool samples. With the increase of intensity of mechanical lysis and integration of lysozyme pretreatment, saturated lysis was achieved by LPQ protocol. And the microbiome structure of samples moved toward one direction when the intensity of lysing condition increases until approaches to saturated lysis. With further investigation, we found that the protocol with saturated lysis condition is more sensitive to detect treatment effects compared to the methods without saturated lysis condition. We hypothesize that the closer a lysing condition of a method to saturation lysis, the more accurate the gut microbiome structure is detected therefore the more sensitive it can be to detect the treatment effect accurately. This indicates that saturated lysing potentially can solve the bias introduced from different DNA extraction methods from different studies and increase the sensitivity of detecting treatment effects.

Therefore, we recommend more DNA extraction methods should be evaluated and optimized using the quality control standards we developed in this study. The standards include increasing the intensity of lysing condition until DNA yield, Gram-positive bacteria ratio, and beta diversity reached a plateau, detecting treatment effects on a same set of samples using methods with non-optimized and optimized lysing conditions. More fecal samples from people with diverse background including people with different diseases should be tested as well. More DNA extraction methods should be evaluated using the same method in this study to confirm the finding of saturated lysing condition increased sensitivity of detecting treatment effects.

## Material and method

### Recruitment, inclusion criteria and exclusion criteria

4 aged between 18 and 35 years old, BMI 20-25, with different ethnicity (White / Caucasian, Black / African American, Hispanic / Latino, and Asian), self-claimed generally healthy participants were recruited in a study approved by Rutgers Arts and Sciences IRB with protocol ID: Pro2018001964. Individuals who are unable to provide the specimen within 90 minutes of sample collection; have any conditions deemed by the investigators that would prevent participation in the study e.g. participation in past or active clinical research, at the discretion of the investigators; have any conditions deemed by the investigators that would compromise the individual’s ability to complete the study, e.g. serious psychiatric conditions, at the discretion of the investigators) were excluded from the study.

5 aged between 18 and 65 years old, have been diagnosed with type 2 diabetes participants were recruited in a study approved by New Brunswick Health Sciences IRB with protocol ID: Pro2019000578. Fecal samples were processed with in vitro fermentation to compare the sensitivity of detecting treatment effects of lysing conditions. Individuals who are unable to provide the specimen within 90 minutes of sample collection, or those who have treatments (currently using or have used antibiotics continuously for > 3 days within 3 months prior to enrollment; currently using or have used weight loss medications or supplements within one month prior to enrollment; use of anti-psychotic drugs), conditions or diseases that are known to alter their gut microbiota (Pregnancy/breast feeding; Self-reported colorectal symptoms or disorders; Diagnosed colorectal diseases or colorectal adenomas; Active or history of malignant tumors; Active or history of liver cirrhosis, chronic or persistent hepatitis; Severe or unstable heart failure; Myocardial infarction within six months prior to enrollment; Any surgery within six months prior to enrollment; Hospital admission for depression; Presence of an eating disorder or purging behavior; Any conditions deemed by the investigators that would prevent participation in the study e.g. participation in past or active clinical research, at the discretion of the investigators; Any conditions deemed by the investigators that would compromise the individual’s ability to complete the study, e.g. serious psychiatric conditions, at the discretion of the investigators) were excluded from the study.

### Homogenized fecal suspension preparation

The samples collected from 4 healthy subjects were stored in box with ice packs surrounded. Fecal samples were transferred to lab within 2 hours of collection. Each fecal sample was homogenized in phosphate buffered saline (pH 7.4, 0.01M phosphate buffer, 0.0027M potassium chloride and 0.137 M sodium chloride, Sigma-Aldrich, Germany) buffer with 1mg/mL resazurin (Sigma-Aldrich, Germany) and 0.05% L-Cysteine hydrochloride (Sigma-Aldrich, Germany) into 25% (w/v) fecal suspension and filtered through cheesecloth. Suspended fecal sample was aliquoted into 1mL aliquots and centrifuged under 16,000 g for 10 min. The supernatant was discarded, and the pellet was stored in -80°C freezer until further use.

### In vitro fermentation

Corn bran (Honeyville, CA, USA), oat bran (GrainMillers, OR, USA), sorghum bran (Nulife, KS, USA), and wheat bran (KSU, USA) were mixed by 1:1:1:1 ratio. The mixture of four brans was blended, roasted at 135°C for 5 minutes, and sifted to make bran mix. Fecal samples collected from 5 type 2 diabetes mellitus patients were stored in box with ice packs surrounded. Fecal samples were transferred to lab within 2 hours of collection. Each fecal sample was homogenized in phosphate buffered saline (pH 7.4, 0.01M phosphate buffer, 0.0027M potassium chloride and 0.137 M sodium chloride, Sigma-Aldrich, Germany) buffer with 1mg/mL resazurin (Sigma-Aldrich, Germany) and 0.05% L-Cysteine hydrochloride (Sigma-Aldrich, Germany) into 25% (w/v) fecal suspension and filtered through cheesecloth inside anaerobic chamber (5% Hydrogen; 5% Carbon dioxide; 90% Nitrogen, Airgas Company, MO, USA). Homogenized fecal suspension for each subject was processed with in vitro fermentation in 3 groups, including 1% Bran Mix (BM), 1% Bran Mix with 1% Inulin (BM+I) and 1% Inulin (IN). The in vitro fermentation was set up in anaerobic chamber (5% Hydrogen; 5% Carbon dioxide; 90% Nitrogen, Airgas Company, MO, USA). Supplements for 3 groups were homogenized with phosphate buffered saline (pH 7.4, 0.01M phosphate buffer, 0.0027M potassium chloride and 0.137 M sodium chloride, Sigma-Aldrich, Germany) buffer with 1mg/mL resazurin (Sigma-Aldrich, Germany) and 0.05% L-Cysteine hydrochloride (Sigma-Aldrich, Germany) to make 1.25x solution. 4mL of supplement solution was mixed with 1mL 25% fecal suspension to get a 5mL culture volume. For each group, triplicates were set up. Three aliquots from each culture were collected and centrifuged. 3 pellets from each culture were used for DNA extractions using 3 methods respectively, including QIAamp PowerFecal DNA kit, PQ, and LPQ.

### DNA extraction using QIAamp PowerFecal DNA kit

DNA was extracted from the pellet of the culture using QIAamp PowerFecal DNA Kit (Qiagen, Germany) according to the manufactory instructions except using TissueLyser II (Qiagen, Germany) at 25 Hz for 10 min instead of vortex at step 5.

### DNA extraction using PQ with 10 lysing conditions

Suspended fecal samples or pellets of in vitro fermentation sample were processed with modified PQ^1^. The modifications include using TissueLyser II (Qiagen, Germany) at 25 Hz instead of Fastprep™ for mechanical lysing step and using nuclease-free water instead of AE buffer to elute the DNA. Step 7 was repeated to increase the rounds of bead-beating. For lysozyme pretreatment, suspended fecal samples were incubated with 100 ul enzymatic lysis buffer (20 mM Tris-Cl, 2mM EDTA, 1.2% Triton X-100, 20 mg/mL lysozyme, Sigma-Aldrich, Germany) at 37°C for 30 min and then processed with modifies PQ with increased bead-beating as above.

### 16S rRNA gene sequencing

Hypervariable region V4 of the 16S rRNA gene was amplified using polymerase chain reaction (PCR) with Ion Torrent barcode tagged primers (515F, 816R) ^11, 12^. PCR reaction was performed using a mixture of 10 µl Platinum™ SuperFi™ DNA Polymerase (ThermoFisher Scientific), 1 µl of 10 µM forward primer, 1 µl of 10 µM reverse primer with unique barcode for each sample, 6 µl of nuclease-free water, and 2 µl of 10 ng/µl extracted DNA solution. The 20 µl reaction mixture will follow the cycling conditions of: 98°C for 30 seconds for initial denaturation; 98°C for 8 seconds, 59.6°C for 10 seconds, 72°C for 10 seconds for 30 cycles of denaturation, annealing and extension; 72°C for 5 min for final extension; hold at 4°C. PCR products were then purified using AMPure XP beads (Beckman Coulter, FL, USA) in 1:1.5 sample to beads ratio to remove primer dimers. All purified amplicons were quantified using Qubit 4 (ThermoFisher Scientific), diluted to 30 pM and pooled into a single library for preparing sequencing chips using Ion Torrent Chef platform. The prepared chip was then sequenced using Ion Torrent S5 platform (ThermoFisher Scientific) following the manufacturer’s protocol.

### Microbiome and Statistical analysis

Primer and adapter removal, denoising and quality filtration of the sequencing data were performed using QIIME 2 to obtain reliable amplicon sequence variants (ASVs) ^13, 14^. Spurious ASVs were further removed by an abundance-filtering method ^15^. A phylogenetic tree was built using the commands alignment mafft, alignment mask, phylogeny fastree, and phylogeny midpoint-root to generate the weighted UniFrac metric. A taxonomy assignment was performed using the q2-feature-classifier plugin based on the sliva database (release 132) ^16^. Sequencing data were then rarefies sampled at 14,400 reads according to rarefaction curve for healthy subjects’ dataset and 14,000 reads for T2DM patients’ dataset. Weighed abundance for each ASV was calculated by multiplying rarefied reads of each ASV and weighted DNA yield. Alpha diversity and beta diversity based on weighted abundance were calculated using QIIME2. All statistical analyses were performed using software R 3.3.2^17^. Weighted DNA yield and Gram-positive bacteria ratio between groups were compared pariwisely using Linear mixed-effects model with subject as random effect and Tukey’s test using R nlme package version 3.1.149^18^ and emmeans package version 1.6.0. Principal Coordinates Analysis (PCoA) and Covariate Adjusted Principal Coordinates Analysis (aPCoA) with subjects adjustment based on Bray Curtis distance were performed to visualize the treatment effect using R VEGAN package version 2.5-6^19^ and R aPCoA package version 1.2^20^. Permutational multivariate analysis of variance (PERMANOVA) with subjects stratified was performed on Bray Curtis distance between groups to test the significance on microbiome structure difference between treatments using R VEGAN package version 2.5-6^19^. Mean Bray Curtis distance within group and between group were calculated and shown as shades in the heatmap. R MaAsLin2 package version 1.2.0^10^ was used for identifying the responses of ASVs toward treatments. The reads of each sample were log transferred and the prevalent ASVs (>20%) were selected for further analysis. Treatment-responsive ASVs were identified by performing generalized linear model for each ASV among samples with different treatments. The p value was then adjusted by the Benjamini-Hochberg procedure. Significant responded ASVs were selected for plotting heatmap. Cladograms were plotted with ward.D2 linkage method using R stats package 4.1.0^21^.

## Acknowledgements

This work was supported by grants from the School of Environmental and Biological Sciences and New Jersey Institute for Food, Nutrition, and Health (seed pilot grant cycle 1, awarded in 2019), Canadian Institute for Advanced Research and Notitia Biotechnologies Company (829441).

## Conflict of interest

Liping Zhao is a co-founder of Notitia Biotechnologies Company.

