## Supplementary Table 1 for "Saturated cell lysing is critical for high sensitivity microbiome analysis"

**Supplementary Table 1 Participants’ information**

|  | **PDM05** | **PDM07** | **PDM09** | **PDM11** |
| --- | --- | --- | --- | --- |
| Sex | F | F | F | M |
| Ethnicity | White | Black | Asian | Hispanic |
| BMI (kg/m^2^) | 20.21 | 21.68 | 24.37 | 22.32 |
| Age | 20 | 21 | 23 | 31 |
